# Hair promotion effect of *Beta vulgaris* L. fruit extracts; physicochemical, and analytical evaluation

**DOI:** 10.64898/2025.12.31.697263

**Authors:** Fareeha Hasan, Siraj Khan, Abeer Tariq, Muhammad Islam

## Abstract

Natural products are used frequently for medicinal purposes due to their lower toxicity effect and greater safety than synthetic products. Alopecia is becoming common due to inheritance, hormonal change and environmental factors. The current study evaluates the physicochemical, analytical and hair growth promotion activity of extracts of *Beta vulgaris* L. Extraction was done by using hexane, chloroform, methanol, ethanol and water in increasing order of their polarity. For the hair growth activity, five groups (positive, negative, untreated, test-group 1 and test-group 2) of albino mice were made and each group has five mice. The hair tonic formulation was applied on their dorsal surface.

The analysis revealed that methanolic and ethanolic extracts showed the highest phytochemical content. The average length and diameter were noted. On day 14 and 21 length of different groups were: standard group (7.40±0.34, 8.15±0.56), negative group (3.05±0.65 and 5.65±0.78), untreated group (3.44±0.27 and 4.58±0.61), test group1 5% methanol (5.25±0.20 and 6.26±0.10), 10% methanol (5.23±0.07 and 6.35±0.33), test group2, 5% ethanol 5.28±0.50 and 6.30±0.84, 10% ethanol 5.50±0.50 and 6.72±0.75 and the diameter of groups were; standard (3.5±0.42 and 4±0.86), negative (1.5±0.33 and 2±0.22), untreated (2±0.34 and 2±0.24), test group1 5% (2±0.41 and 2.5±0.65), 10%(2±0.65 and 2.5±0.34), test group2 5%(1.5±0.67 and 2.5±0.42), 10% (2±0.88 and 2.5±0.54). Both extracts showed significant results, but ethanolic extract is more effective than methanolic. The exact mechanism of hair growth is not known yet, but it might be due to its strong antioxidant and anti-inflammatory effect. hence, this study is useful in treating baldness.

## 1. Introduction

Nature is paradigm of synergism. Due to harmful effects of synthetic drugs, there is escalating heed towards nature for the development of new drugs. Approximately 80% of the population of developing countries hinge on conventional medicine attained from plants and animals for their ultimate well-being. In the US around 25% of prescriptions have a minimal one naturally attained active ingredient. Different primordial systems like Ayurvedic, Unani, Siddha and Allopathy use herbs for several maladies (Caesar and Cech, 2019). The therapeutically active compounds from old times are excellent source for rational drug design. It would be more fruitful and least expensive to reconsider the data of traditional herbal treatments from earliest times (Chaachouay and Zidane, 2024).

Herbal medicines in Asia have a long history of human interactions with human surroundings. Plants used in traditional medicine have a variety of ingredients that can be used for short and long-term treatment (Liu, 2021). Heeding towards plants shows sustainability of conventional claims of beneficial effects of natural products for human care (Pan et al., 2014). Conventional medicine is the implementation of various thoughts, beliefs, experiments originating from different cultures and times. Leading to cure diseases, improving health of mankind (Che et al., 2024).

Human history is antedated with use of plants for curing purposes and still plants are source of current medicines and are highly effective in their actions for example quinine (from cinchona bark), digoxin (from foxglove), morphine (from opium poppy), and aspirin (from willow bark). On a big scale in collaboration with drug companies’ plants are being scrutinized for their pharmacological actions and leading to drug development (Claro et al., 2024; Petrovska, 2012). To look out for new novel drugs which must be efficacious enough to combat with developing diseases. From thousands of years traditional medicines have been used to treat different diseases and now a days these are viable source for the treatment due to increased drug resistance and affordability for example antimalarial drugs (Willcox and Bodeker, 2004).

Hair is an important part of the human body and gives an overall attractive appearance (Dhami, 2021). They come from the upper layer of skin. Hair plays a protective role and is considered equivalent to sweat glands, nails and sebaceous glands. They are epidermal offspring because they have developed from the epidermis in the embryo (Kaushik et al., 2011). Hair is protective against environmental influences like UV light and temperature (Ji et al., 2021). There are many problems with hair like depigmentation, hair loss and dandruff. Continuous hair loss ultimately causes baldness and lower hair volume (Nandaniya et al., 2023). Approximately 100,000 to 150,000 hairs are present out of which 50 to 100 strands sheds normally (Li et al., 2022). Cyclic hair growth is important to prevent hair loss in which a strand of hair is formed that grows from a follicle and detaches and a new hair grows from the same follicle (Natarelli et al., 2023).

Herbal medicine has been proven to be effective in treating many diseases. Beta vulgaris is a plant that is native to the Mediterranean, on the Atlantic coast of Europe, the Middle East and India (Miraj, 2016). They are a biennial herbaceous, occasionally annual crop, cultivated for their edible roots and leaves (Mirmiran et al., 2020).

The roots and leaves of beets have been used in folk medicine to treat a variety of diseases. It is used as a laxative, for wound healing, as an aphrodisiac, for digestion and for blood disorders. “Garlic breath neutralizing effect” against oxidative stress, neuroprotective effect, antifungal, antihyperglycemic, anti-inflammatory, cancer-fighting effect (Miraj, 2016). Red beet is used in food, paintings, ornamental art. Medicinally used to combat many common diseases related to blood, heart, liver, pancreases, digestive and neurological (Kapadia and Rao, 2013). Traditionally its juice is used for sexual dysfunction, for removal of kidney stones. Hipprocrates used their leaves to alleviate wounds (El Gamal et al., 2014). Red beet (Beta vulgaris) juice (RBJ) is used as a food item, as a traditional medicine for cosmetics and for the treatment of anemia (Babarikins et al., 2013).

Red beet possess good antibacterial properties due to the presence of phytochemicals and can be incorporated in different formulations for human health like antimicrobial food packaging (Manohar et al., 2017), cosmetics like petroleum jelly, lotions and oils, drops, creams, soaps, shampoos, toothpaste, deodorants, and hair dyes. Hence not only impart color but also prevent aging related effects. Also, dietary supplements, topical sprays, and ointments targeting diseases (Kumar and Brooks, 2018).

In skin care products it shows promising effects acting as natural detoxifier, preventing inflammation, resulting in radiant skin. It treats acne, for example *S. aureus* bacteria causing acne is highly susceptible to red beet extract *in vitro* (Bezalwar Pratik et al., 2014; Clifford et al., 2015; Martinez et al., 2015).

Beets are rich in nitrogenous compounds and phytochemicals which play many important roles in human health like they are useful for thrombophlebitis, varicose veins, hypertension largely effect systolic blood pressure, increasing stamina and reduces demand of oxygen required during exercise and other things. It removes bad cholesterol from the body, natural detoxifier for the body (Maheshwari et al., 2013). Hence the use of red beet is preventive against age-related diseases (Singh and Hathan, 2014). For example, endothelial dysfunction (important for maintaining vasoprotective functions, which otherwise increases risk for atherosclerosis, cardiovascular functions etc), cognitive impairment (Clifford et al., 2015).

## 2. Materials and Methods

### 2.1 Plant Material

Red beet root *Beta vulgaris L.* was collected from the local market of Lahore Punjab Pakistan. Beet root was identified and authenticated from Department of Botany, Government College University, Lahore by Prof. Dr Zaheer-ud-din Khan. *Beta vulgaris L.* was deposited in the herbarium of the GCU (GC. Herb. Bot. 3664). The leaves of beetroot were removed. Root was peeled off, sliced and shade dried. Collected, then sieved to remove any dust or external particle and stored at room temperature.

### 2.2 Chemicals and Solvents

Analytical grade quality of solvents and chemicals were used. The names and sources of chemicals are given below:

Acids: Concentrated hydrochloric acid (BDH, England), concentrated nitric acid, concentrated sulfuric acid, organic solvents like acetone (E. Merck A.G Damstadt, Germany), aluminium nitrate, anthrone reagent (Sigma life sciences, Germany), bovine serum albumin (Bioshop, Canada), chloroform, copper sulfate, ethanol (BDH, England), Foiln-Ciocalteu’s reagent, gallic acid (Sinochem, China), glucose, n-hexane, methanol, petroleum ether, potassium acetate (Peking chemical works, China), quercetin (Merck, Germany), sodium hydroxide, sodium carbonate, sodium potassium tartrate, and triton-X (Unichemicals, Ireland). (Daejung, China).

### 2.3 Extraction

Extraction is a procedure to isolate the required compounds by using particular solvents for extraction. We employed two types of extraction techniques in this study which were: hot extraction and cold extraction.

#### 2.3.1 Hot Extraction

Powder of beet root weighing sixty grams was subjected to serial extraction with organic solvents. The solvents were used according to their increasing order of polarity, firstly n-hexane (B.P 42°-62°C), then Chloroform (B.P 61°-62°C) and finally with Methanol (B.P 54°-64°C). The weighed powder was poured on filter paper to make it thimble, which was then placed in thimble assembly of soxhlet. Then 500ml of the solvent n-hexane was added to flask of apparatus. Extraction was carried out until the siphon was cleared at temperature of 40°C. Solvent was collected after extraction and put in rotary evaporator to concentrate at temperature below the boiling point of the solvent. The concentrated extract was then shifted to airtight vials, after that it was placed in oven to dry completely at 25°C. This process was performed again with the other two solvents using the same thimble. The percentage yield was calculated in accordance with the amount of powder taken initially (Yadav and Agarwala, 2011).

#### 2.3.2 Cold Extraction

In 1000ml beaker twenty-five gram of beet root powder was taken and 250ml of ethanol was added in the powdered material. It was stirred for 24 hours at 50 revolutions per minute with the magnetic stirrer. The filtrate was collected after filtration and to the residue another 250ml of ethanol was added and subjected to stirring with magnetic stirrer for one hour. This procedure was repeated twice, and filtrate was collected separately. The extract was then subjected to dry in rotary evaporator to semi-solid state. For aqueous extraction the same technique was employed, and the resulted extract was dried in rotary evaporator at 60°-65°C which is bridged to ultra-low chiller of temperature 4°C (Azwanida, 2015).

### 2.4 Physicochemical Analysis

According to USP 2005 methods, physicochemical analysis of powdered material of red beet root *Beta vulgaris* was performed.

#### 2.4.1 Moisture Content

Taken 2 grams of accurately weighed powder in silica crucible, which was previously washed, dried and weighed by using electronic balance. The weight of empty crucible was noted. After that powder was put in oven for drying at 105°C for 30 minutes. The weight of crucible along with powder was noted after every 30minutes until the weight becomes constant. To cool down the crucible to room temperature it was put in desiccator. The weight loss in powder was determined and by using the following formula moisture content was calculated.

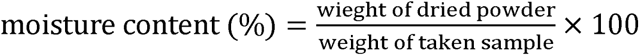

#### 2.4.2 Ash Value

The ash values were determined according to methods described in USP (2005). Ash values include total ash, sulphated ash, water soluble ash and acid insoluble ash.

#### 2.4.3 Total Ash

Two grams of powder were taken in clean, dry, and weighed silica crucible. Then it was incinerated in muffle furnace whose temperature was raised gradually to 675±25°C until powder becomes carbon free (white in color). The crucible was then allowed to cool in desiccator to room temperature. The total ash percentage was calculated by using the formula given below:

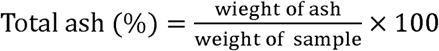

#### 2.3.4 Acid Insoluble Ash

The total ash obtained by the above-mentioned method was boiled in 25ml of 3N HCl for 5 minutes and cooled down at room temperature. By using less ash filter paper, the insoluble matter was filtered and the obtained residue over filter paper was washed with hot distilled water. Then it was placed in weighed silica crucible and put in muffle furnace. Heated until it became carbon free and crucible was allowed to cool at room temperature and weighed afterwards. From the air-dried material, the percentage of acid insoluble ash was calculated.

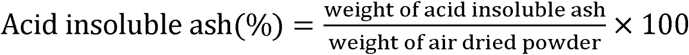

#### 2.3.5 Water Soluble Ash

The obtained total ash from above procedure was allowed to boil with 25ml of distilled water for 5 minutes. The insoluble water substance was then filtered using ash less filter paper. The retained residue was washed with hot distilled water. The filter paper along with residue was put in previously weighed silica crucible. It was placed in a muffle furnace for incineration for about 15 minutes at temperature not raising than 450°C, till it becomes carbon free. By using the given formula, the percentage was determined.

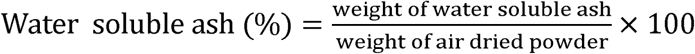

#### 2.3.6 Sulphated Ash

In clean, dry and weighed silica crucible, two grams of powder were taken, to which 1ml of concentrated sulphuric acid was added. It was then gently heated on flames till no white fumes formed. This process was repeated twice. For 30 minutes the crucible was placed in muffle furnace for incineration at 675 ± 25°C and cooled to room temperature. For percentage calculation, the silica crucible was weighed after cooling and following formula was applied.

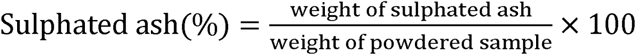

#### 2.3.7 Extractive Values

The extractive values include aqueous soluble extractive value and alcohol soluble extractive value, which was determined by established procedure (Arawande et al., 2018).

#### 2.3.8 Alcohol Soluble Extractive

Five grams of powder was taken to glass stoppered conical flask and for 24 hours macerated with 100ml of 90% of ethanol, with efficacious stirring. This blended material was then filtrated. To weighed China-dish, add 25ml of the filtrate and evaporate it to dryness at 105°C until constant weight is achieved. The extractive value was calculated with reference to dried sample powder by using the given formula below:

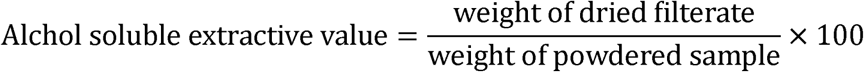

#### 2.3.9 Water Soluble Extractive

To glass stoppered conical flask five grams of powder was taken. For maceration add 100ml of chloroform water (which is prepared by mixing 30ml of chloroform with 70ml of distilled water) for 24 hours with effectual agitation. Afterwards mixture was filtrated and to previously weighed China-dish add 25ml of the filtrate, which was evaporated to dryness at 105°C in oven to the constant weight. The extractive value was calculated with reference to dried sample powder by using the given formula below:

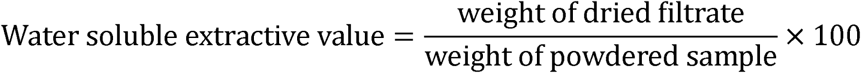

#### 2.3.10 Estimation of Primary Metabolites

Estimation of primary metabolites present in powdered plant material includes total protein contents, total lipid contents and total carbohydrates.

#### 2.3.11 Total Proteins Content

The evaluation of total protein content was determined using method given (Lowry et al., 1951). To test tube, add 500 mg of powdered plant material, with 10ml distilled water in which two-three drops of triton-X were added. The mixture was incubated for 30 minutes with continuous agitation, afterwards the mixture was centrifuged for 10-15 minutes at 2700 revolutions per minute. The 100 µl of supernatant layer was taken in a test tube and volume make up to 1ml was done with distilled water. To this test tube add Folin-Ciocalteau reagent (200µl) and 3ml of Reagent C which is made from 50ml of reagent A (prepared by mixing 2% sodium carbonate in 0.1N sodium hydroxide) and 1ml of reagent B (prepared from 0.5% copper sulfate in 1% potassium sodium tartrate). Left the mixture for 30 minutes incubation at room temperature. As a standard Bovine serum albumin (BSA) aqueous solution (1mg/ml) was used. BSA standard was formulated in 12.50-100 µg/ml sequence concentrations to plot a calibration curve. Preparation of blank solution involves all the reagents except sample. Absorbance of all prepared solutions was checked at 600nm against blank. Total proteins content was then calculated from the standard curve by linear regression equation.

#### 2.3.12 Total Lipids Content

The method for the evaluation of total lipids was reported (Besbes et al., 2004). Sixty grams of powder material of plant *Beta vulgaris* L. in packed thimble was subjected to soxhlet apparatus. For extraction hexane was used as solvent. After extraction, the extract was dried using rotary evaporator at 40°C. The dried weight of extract was noted, and total lipids content was determined.

#### 2.3.13 Total Carbohydrates

The total carbohydrates content was determined by using (Al-Hooti et al., 1998)

formula: Total carbohydrates (%) = 100 – (total moisture + total ash + total fat +total proteins)

### 2.4 Estimation of Secondary Metabolites

#### 2.4.1 Total Polyphenols

Total polyphenols were evaluated by method described by (Meda et al., 2005).As a standard to gallic acid was used to make the standard calibration curve. The stock solutions of standard gallic acid and samples were made in methanol having concentration 1mg/ml. The further dilutions of both sample and standard were prepared by diluting 10, 20, 40, 80 and 120µl of stock solution up to 1ml with methanol. Take 200 µl of samples and standard solutions in tubes and add 200µl of Folin-Ciocalteu reagent (0.2ml FC reagent) to these tubes and mixed gently. Now add 1ml of 15% Na2CO3 to these test tubes after 4 minutes and make-up final volume to 3ml with methanol. Blank was prepared by the same method as that of sample except the addition of sample which is replaced by 200µl methanol and then all the solutions were incubated for 2 hours at room temperature. The absorbance of solutions was measured at 760nm by using spectrophotometer. Total polyphenol contents were evaluated by linear regression equation from standard curve.

#### 2.4.2 Total Flavonoids

The total flavonoids were evaluated by method given by (Pallab et al., 2013).Quercetin was a standard. Standard stock solution of quercetin and extracts of plant were prepared in methanol of concentration of 1mg/ml. Further dilutions of 10, 20, 40, 80 and 120µl of standard stock solutions were prepared in methanol. Taken 200µl of sample solutions and 200µl standard solutions in test tubes and reagents were added 100µl each of 10% aluminum nitrate solution, 1M potassium acetate and 4.6 ml of distilled water. After that all the solutions were incubated at room temperature for 45 minutes. Blank was prepared same except the addition of sample. The absorbance was measured against blank at 415nm by using a spectrophotometer. The flavonoid contents were evaluated by linear regression equation from the calibration curve.

#### 2.4.3 Total Polysaccharides

The total polysaccharides were estimated by (Abubakar et al., 2015) described method. Each 200mg extract was taken for sample solutions in 7ml of 80% ethanol in centrifuge tubes to remove soluble sugars. It was mixed for 2 minutes on vortex mixer, then tubes were centrifuged for 10 minutes, and supernatant was collected. Similar procedure was repeated until residue didn’t give any color with anthrone reagent (200mg per 100ml H2SO4). Collect all the supernatant in 100ml volumetric flask. The final residue obtained was dried in the water bath and then added 10ml of distilled water and 10 ml of HCl (25%v/v) mixture in 1:1 in it. The mixture was kept for 20 minutes at temperature 0°C in an ice bath and then all tubes were centrifuged for 10 minutes, and supernatant was saved. Extraction was done twice, and all collected supernatant was pooled together in volumetric flask and made-up final volume to 100ml with distilled water. 0.1ml (100ul) of the supernatant was transferred to test tube, made volume with 1ml of distilled water and added 4ml of anthrone reagent. The mixture was heated in water bath at 100°C for 8 minutes and cooled rapidly and the intensity of green color produced was measured at 630nm. 1ml of distilled water and 4ml of anthrone reagent was added in test tube to prepare blank solutions. Glucose was taken as standard. Standard stock solution was prepared in 1mg/ml concentration and further dilutions of 20, 40, 60, 100, and 200µg/ml were prepared. 100µl from above each of the dilutions was added in test tubes and then made final volume with 1ml with distilled water and 4ml of anthrone reagent and the absorbance measured at 600nm.The polysaccharides concentration was evaluated from the standard curve from the linear regression equation and multiple with the correction factor 0.9.

#### 2.4.4 Total Glycosaponins

Total glycosponins were determined by the method described by (Saleem et al., 2014). For this method, about 1g of each of the extracts of *Beta vulgaris* red beet root *was* refluxed for about 30 minutes in 50 ml of methanol. The process was repeated, and extracts were pooled together. The extracts were then concentrated by using the rotary evaporator until the final volume becomes 10 ml. The final amount of extracts was added drop wise in weighed beaker containing 50 ml of acetone. The precipitate obtained was dried in oven at 100°C and weighed to calculate the total glycosaponins value.

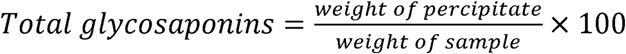

### 2.5 Analytical Studies

#### 2.5.1 Fourier Transform Infrared (FTIR) Scanning

The tip of the sample press knob was elevated slightly from the diamond sampling window. A small amount of material to be analyzed was placed on diamond crystal. It was made sure that the sample covered the surface area of the diamond crystal. Spectrum was then scanned in the range of 4000-400 cm^−1^.

#### 2.5.2 Mineral Content Analysis

By atomic absorption spectrophotometer (PG-990, China), the mineral constitutes of *Beta vulgaris* were determined by method given by(Object, n.d.). Approximately 1 g of dried and properly weighed sample was taken in crucible and then it was put in furnace at temperature of 550°C for 1 hour.

In the hot solution of 10 % HCl and HNO_3_ (ratio 3:1), the ash was dissolved and with distilled water final volume was made upto 100ml with distilled. After that mineral contents were evaluated by using atomic absorption spectrophotometer.

#### 2.5.3 Ultraviolet-Visible profiling

Prepare the stock solution in methanol with concentration of 1mg/ml for each of the extracts. Then 500µl was taken from stock solution and diluted to 10 ml by adding methanol. As blank methanol was used.

Each of the solutions was then scanned in UV-Visible region having range of 200-800 nm and max was recorded.

### 2.6 Experimental animals

Twenty-five, Albino mice, weighing 20±10 grams, were purchased from animal house of Veterinary Research Institute, Lahore, Pakistan. In animal house of University College of Pharmacy, University of the Punjab, Lahore, Pakistan, mice were accustomed for one week, in air-conditioned room 24±2°C with light and dark period of 12h/12h. Mice tail were marked using permanent marker and kept in cages. All the cages have wood shavings to absorb feces, and they were cleaned every third day with disinfectants to maintain the hygiene. Mice were fed with the standard pellet diets and were freely permitted to drink water from water bottles(Sabarwal et al., 2009).

#### Ethical approval

The research design was ratified by Animal Ethical Committee of Department of Zoology, University of the Punjab, Lahore, Pakistan, under vide reference No.AEC.679.

### 2.7 Hair Tonic Formulation

All the components of the formulation have been given in Table 1 below. Methanol and ethanol extracts of *Beta vulgaris* were diluted with distilled water and a good mix was given with help of vortex mixer. To solution of extracts, 98% Ethanol was added and by ultra-sonic mixer it was homogenized. Lastly, butylene glycol was added to the tonic formulation (Imtiaz et al., 2017).

**Table 1:**
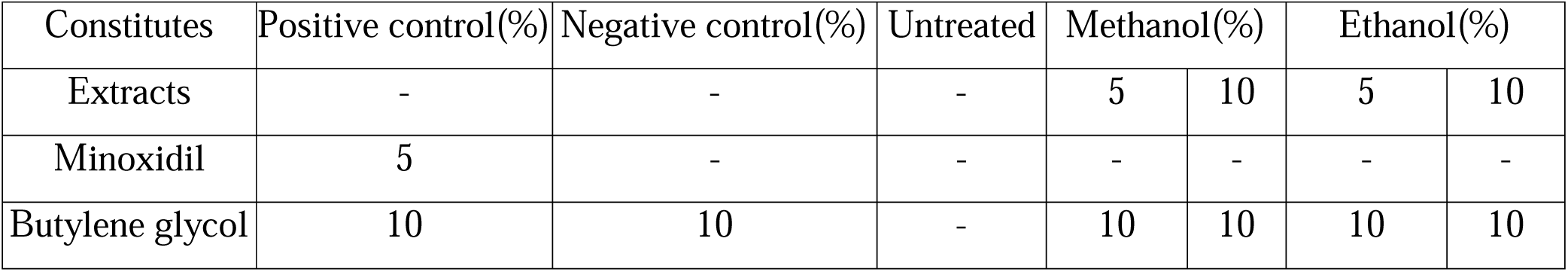
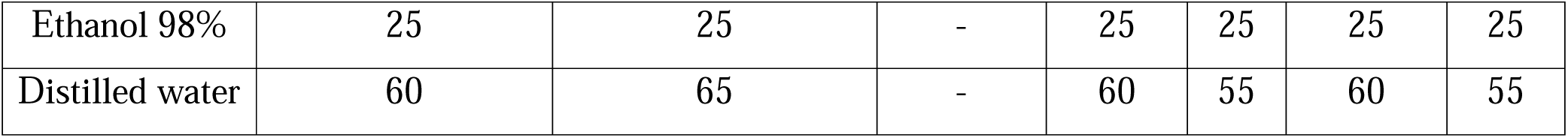
tonic formulation.

#### 2.7.1 Evaluation of Hair Tonic

Assessment of hair tonic was done by following test, 1) organoleptic test: tonic smell and color was noticed, 2) homogeneity test: hair tonic about 100μl was taken on glass dish, smeared and inspected under light to check any coarse particle, 3) pH test: tonic pH was checked by pH meter, which was calibrated by utilizing buffer solution of pH 4 and 7 (Imtiaz et al., 2017).

#### 2.7.2 Hair growth activity

Twenty-five mice of either sex was taken and divided in five groups as positive control, negative control, untreated, methanol and ethanol. In positive control group mice were treated with standard minoxidil 5% solution, negative group was treated with solution having no active ingredient, untreated group (no treatment was given), and two test groups which includes two different extracts that are methanol extract and ethanol extracts. The test group was further divided into two different concentrations of 5% and 10% solutions. Hair of mice was removed from the dorsal side by using hair erasing cream VEET® cream. The shaved part was first rinsed and then cleaned with antiseptic like ethanol. After hair removal mice were observed for 24hours for any sensitivity reaction of the hair removal cream. Before starting the activity mice dorsal surface was divided into two halves left and right parts for the application of the test drugs on it. After the safe observatory period of 24 hours, 100μl of each of the formulation of hair tonic was applied onto the dorsal area of mice for 21 days, twice a time daily. The day one of application of the formulation was considered zero (Imtiaz et al., 2017).

#### 2.7.3 Qualitative hair growth estimation

Qualitative evaluation of hair growth was done by the visual inspection of the two parameters, 1) initiation time of hair growth: that is minimum time required for beginning of hair growth on shaved skin and 2) completion time of hair growth: minimum time required to fully cover the shaved area with new hairs (Sabarwal et al., 2009).

#### 2.7.4 Determination of hair length

From the dorsal side of mice about ten hair strands were randomly tugged on 7, 14 and 21 day. The hair strand was then straightened out and by using digital Vernier caliper, the length was determined. To examine the significance among the groups, the mean length was determined (Adhirajan et al., 2003).

#### 2.7.5 Determination of hair diameter

The diameter of hair was determined by utilizing microscope with ocular micrometer. Statistically the mean diameter was determined (Imtiaz et al., 2017).

## 3. Results and Discussion

### 3.1 Percentage Yield of Extracts

Choice of organic solvents has a vital role in the percentage yield of extracts (Dhawan and Gupta, 2017). Table 2 shows percentage yields of different extracts of *Beta vulgaris*. The maximum percentage yield is methanol, which shows that methanol is the most effective one to extract the components.

**Table 2.** Yield of extracts of root of *Beta vulgaris*.

| No. | Extracts | Percentage yield (w/w) |
| --- | --- | --- |
| 1 | Hexane | 1.15% |
| 2 | Chloroform | 0.9% |
| 3 | Methanol | 50.86% |
| 4 | Ethanol | 12.17% |
| 5 | Water | 10.09% |

### 3.2 Physicochemical Analysis

Physicochemical analysis was studied on root of *Beta vulgaris*. The results are given in Table 3. They were 5.80±0.03 moisture content, 8.31±0.05 total ash, 58.23±0.05 acid insoluble ash, 44.57±0.04 water soluble ash, 34.25±0.03 sulphated ash, 15.0±0.01 alcohol extractive value and 16.88±0.02 water soluble extractive value. Physiochemical analysis of powder of root of *Beta vulgaris* L.

**Table 3.** Physiochemical analysis of powdered root of *Beta vulgaris* L.

| No. | Physicochemical Properties | Percentage Contents ± S.D |
| --- | --- | --- |
| 1 | Moisture content | $5.80 \pm 0.03$ |
| 2 | Total ash | $8.31 \pm 0.05$ |
| 3 | Acid insoluble ash | $58.23 \pm 0.05$ |
| 4 | Water soluble ash | 44.57±0.04 |
| 5 | Sulphated ash | 34.25±0.03 |
| 6 | Alcohol soluble extractive | 15.0±0.01 |
| 7 | Water soluble extractive | 16.88±0.02 |

Total ash value shows the clarity of crude drug by assessing different impurities in it, for example silicate, oxalate, phosphates and carbonates. Higher ash values indicate that certain plant materials have large deposits of minerals in them. Water soluble ash is used to calculate the inorganic components while acid insoluble ash is used for silica and earthy type material. Sulphated ash shows the existence of volatile inorganic compounds. Moisture contents estimate the durability and prevention against degradation process of components of drug during period of storage. Lower values of moisture content show lesser chance of degradation. Prevents microbial growth and enzymatic changes. Important for preservation. Extractive values are used to calculate certain amount of bioactive components in given samples when extracted with specific solvent (Chanda, 2014; Essiett and Bala, 2011; Guiné and A. M. Castro, 2003).

### 3.3 Estimation of Primary Metabolites

The percentage contents of primary metabolites of *Beta vulgaris* L. were assessed and given in Table 4.

**Table 4.** Estimation of primary metabolites of powder of *Beta vulgaris* L.

| No. | Primary Metabolites | Percentage Contents ±SD |
| --- | --- | --- |
| 1 | Total proteins | 1.35 ± 0.004 |
| 2 | Total lipids | 1.15 ± 0.002 |
| 3 | Total carbohydrates | 83.39 ± 0.001 |

Primary metabolites possess basic nutritional importance in plants. Higher percentage of carbohydrates indicates that *Beta vulgaris* is power pack of energy while lesser value of lipids estimates that it has no or lower cholesterol levels.

### 3.4 Estimation of Secondary Metabolites

Total flavonoids, total polyphenols, total polysaccharides and total glycosaponins are secondary metabolites which were analyzed, and results are given in Table 5.Calculated linear regression equations were total polysaccharides (y = 0.0055x - 0.0163, R^2^ = 0.9945), total flavonoids (y = 0.0046x - 0.0008, R^2^= 0.9971), total polyphenols (y = 0.001x - 0.0059, R^2^ = 0.9966).

**Table 5.**
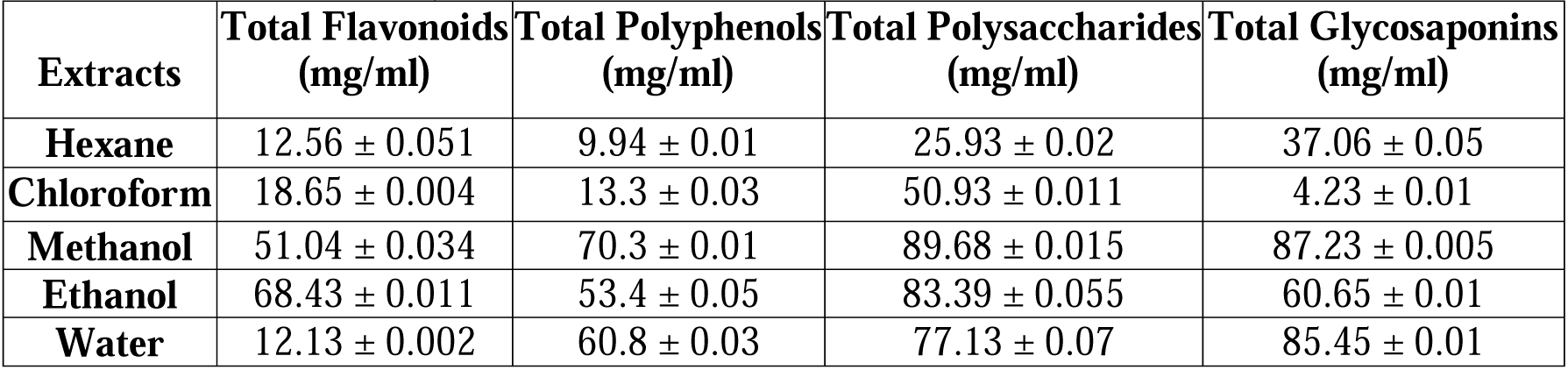
Secondary metabolites of different extracts of *Beta vulgaris* L.

| Extracts | Total Flavonoids<br>(mg/ml) | Total Polyphenols<br>(mg/ml) | Total Polysaccharides<br>(mg/ml) | Total Glycosaponins<br>(mg/ml) |
| --- | --- | --- | --- | --- |
| Hexane | 12.56 ± 0.051 | 9.94 ± 0.01 | 25.93 ± 0.02 | 37.06 ± 0.05 |
| Chloroform | 18.65 ± 0.004 | 13.3 ± 0.03 | 50.93 ± 0.011 | 4.23 ± 0.01 |
| Methanol | 51.04 ± 0.034 | 70.3 ± 0.01 | 89.68 ± 0.015 | 87.23 ± 0.005 |
| Ethanol | 68.43 ± 0.011 | 53.4 ± 0.05 | 83.39 ± 0.055 | 60.65 ± 0.01 |
| Water | 12.13 ± 0.002 | 60.8 ± 0.03 | 77.13 ± 0.07 | 85.45 ± 0.01 |

Saponins are used as major components for drug formation from plants and folk medicinal products. It has important pharmacological functions such as anti-inflammatory, anti-fungal, cytotoxic, anti-tumor, anti-viral, anti-parasitic activities (Sparg et al., 2004). Methanolic extract has the highest glycosaponins contents as saponins are more soluble in polar organic solvents (Shi et al., 2004).

Polyasaccharide has excellent properties of biodegradability and biocompatibility, so achieving good polymer property to be used as biomaterial for example anti-tumor, anti-viral and anti-bacterial (Lemarchand et al., 2004). The percentage of polysaccharides is higher in methanolic extract of *Beta vulgaris*.

The percentage of total polyphenols is high in methanol extract whereas for total flavonoids ethanol extract contains maximum concentration. Polyphenols are multiple phenolic compounds found in food, possessing excellent antioxidant properties. They are useful as anti-cancer, anti-tumors, cardioprotective, neurodegenerative diseases and treating diabetes milletus (Scalbert et al., 2005). Flavonoids are biologically active polyphenolic compounds, which have great anti-oxidant activity. Possess anti-inflammatory, anti-hepatotoxicity activities and various other health benefits (Chhikara et al., 2019; Rose et al., 2014).

### 3.5 Fourier Transform Infra-Red Spectroscopy Studies

FTIR evaluation shows the presence of various functional groups such as alkanes, alkenes, aldehydes, ketones, polysaccharide, aromatic, carboxylic acid, phenols, amines or nitro compounds, alkyl halides or halogen compounds. At range of 3650-3200 cm^−1^, all extracts have strong and broad peak for O-H (alcohol) bond stretching while at 2500-3300 cm □^1^ O-H (carboxylic acid) bond stretching. At range of 3000-2700 cm^−1^ have strong and sharp peak for C-H bonds stretching whereas at 3300-3500 cm □^1^, the chloroform and ethanol extract have strong and broad peak for N-H stretching bonds. The extracts of hexane and chloroform, indicates the presence of C=O stretching bond at 1741.72 cm □^1^ and 1712.72 cm □^1^ respectively. The methanol extract showed presence of diketones C=C at 1634 cm□^1^ while at range of 1550-1650 cm□^1^ and 1600-1680 cm□^1^ indicates C=N and C=C stretching of bonds respectively, in ethanol, water and hexane extracts. The methanol, ethanol and water extracts showed the presence of alkanes and O-H bending bonds at 1408cm^−1^, 1406 cm□^1^, 1408 cm□^1^ respectively while at 1515.95 cm□^1^ chloroform extract indicates alkanes and alkyl groups. The chloroform extract at 1455.59 cm□^1^ shows nitrosamine group, at 1366 cm□^1^ shows isopropyl group and 1258 cm□^1^ shows ester carbonyl presence. The range of 1230-1020 cm□^1^ and 1250-1050 cm□^1^ specifies C-N and C-O bonds presence severally, in extracts of methanol, ethanol, water and hexane. It shows that it contains alkyl amines and alkyl ketones. The aromatic compounds are present in methanol and water extract at 830 cm□^1^ and 831 cm□^1^ respectively. Peaks at range of 500-730 cm□^1^ shows chloro halogen compound and 490-620 cm□^1^ shows iodo halogen compound. All extracts show existence of chloro halogen compounds whereas the extract of methanol and water also shows iodo halogen compounds (Chen et al., 2001; El-Hendawy, 2006; Ovchinnikov et al., 2016; Petit and Puskar, 2018; Sim et al., 2004). Presence of various functional groups in extracts specifies medicinal value of root of *Beta vulgaris* L. FTIR Scan of different extracts of root of *Beta vulgaris L. are given in Figure 1*.

**Figure 1.**
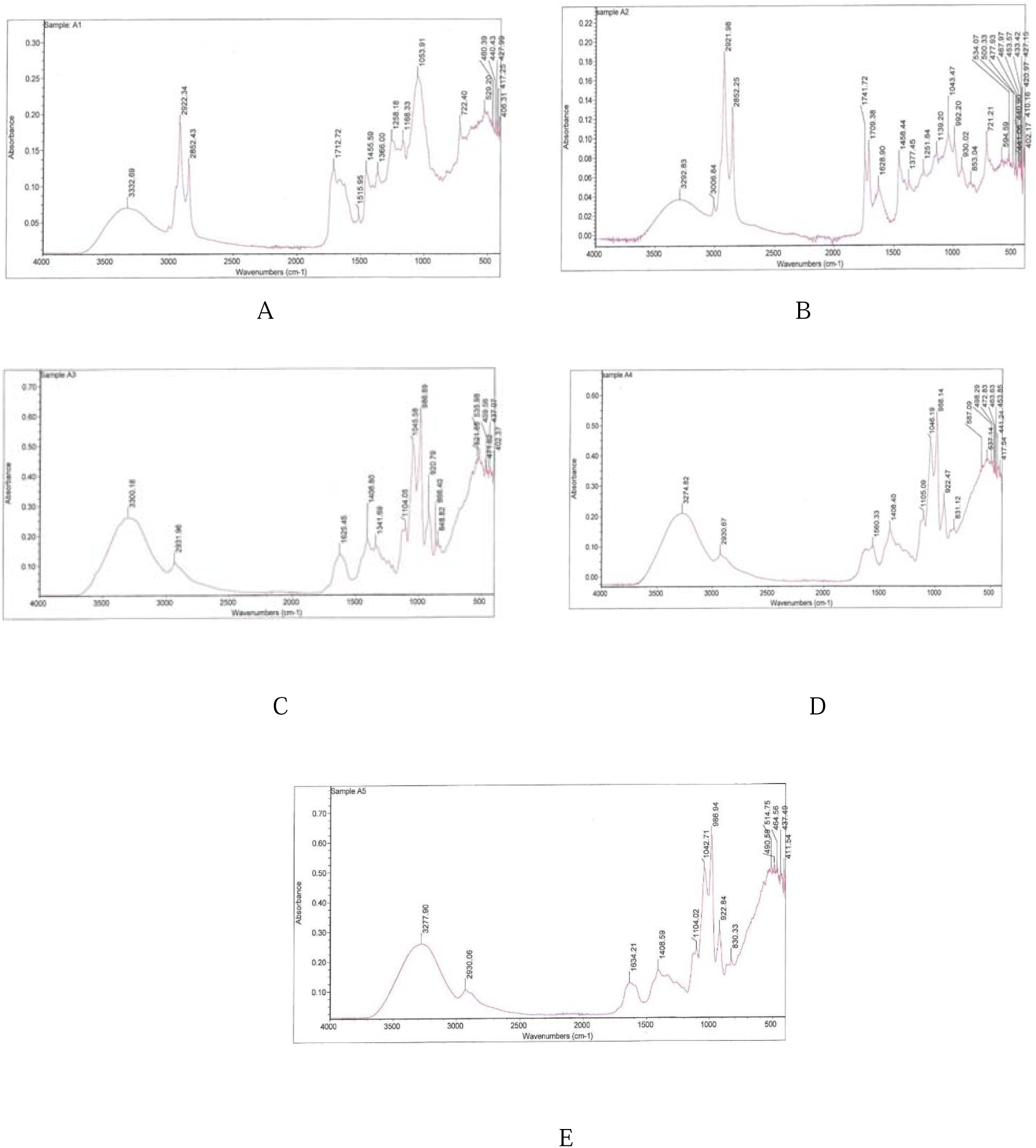
FTIR scan of (A) Chloroform, (B). hexane, (C). ethanol, (D). water, and (E). methanol extract of the root of *Beta vulgaris* L.

### 3.6 Estimation of Mineral contents using Atomic Absorbance Spectrophotometer

The powder root of *Beta vulgaris* was subjected to test minerals like sodium, magnesium, potassium, calcium, phosphorous, iron and zinc by using atomic absorbance spectrophotometer and results are given below in Table 6

**Table 6.** Mineral estimation of powder of root of Beta vulgaris L.

| Elements | Concentration (mg/ml) |
| --- | --- |
| Calcium | 0.16 |
| Magnesium | 0.23 |
| Sodium | 0.78 |
| Potassium | 3.04 |
| Iron | 0.0079 |
| Zinc | 0.0037 |
| Phosphorous | 0.39 |

The result indicates that the root of *Beta vulgaris* L. contains high concentration of potassium. The concentration of sodium, phosphorous, magnesium and calcium are lower. It also contained iron and zinc in minute quantities. Minerals are essential for human body for example potassium is necessary for nerve impulses transmission, for contractions for smooth, skeleton and cardiac muscles, for maintaining blood pressure and it’s also good for bones and useful for protein synthesis. Out of total body calcium, majorly present in teeth and bones. Important for normal rhythm of heart, secretion of hormones, clotting of blood, activation of enzymes and contraction of muscles. Magnesium is a major marco-nutrient useful in biological processes like protein and nucleic acid metabolism, as a co-factor in many enzymes, in neuromuscular transmission, for contraction of muscles, for growth of bones and for regulation of the blood pressure. It is important for the metabolism of calcium and for potassium refluxes. Sodium and potassium play an essential role in maintaining ionic balance in the body. Sodium is essential in maintenance of osmotic pressure, acid-base balance, membrane potential and also useful for active transport. Phosphorous is necessary for many physiological activities in the human body, it occurs as important biological component like in lipids, carbohydrate, proteins and nucleic acids. In bone formation, buffers and in sugar metabolism. Iron and zinc are essential for energy production and for enzymatic anti-oxidants. Iron is vital for utilization, transportation and storage for oxygen. It is an important component of hemoglobin, myoglobin, cytochrome and other types of proteins. Zinc is important for the immunity of human body, for growth, sexual development, it assists in many physiological and metabolic process, and wounds healing (Babarykin et al., 2019) (Klevay and Combs Jr, 2005; Mir-Marques et al., 2016; Rop et al., 2012; Soetan et al., 2010; Zamberlin et al., 2012)

### 3.7 Ultra-Violet Visible Profiling

UV-VIS spectroscopy is useful quantitative technique for determination of organic or inorganic components present in various samples, either in solution or gas form (Mohammed, 2018; Zagatto and Worsfold, 2017). Figure 2 shows the results of extracts of *Beta vulagris* L. after UV-Vis analysis. All the extracts showed the absorption in near ultra-violet range of 180-390nm. Hence, the same absorption pattern for all of them. In this range showed presence of certain phenolic components like flavonoids. As, some of the Flavonoids lie in the range of 250-300nm (Sisa et al., 2010). Along with the UV range, they also fall in visible range too (Aleixandre-Tudo et al., 2018; Julkunen-Tiitto et al., 2015).

**Figure 2.**
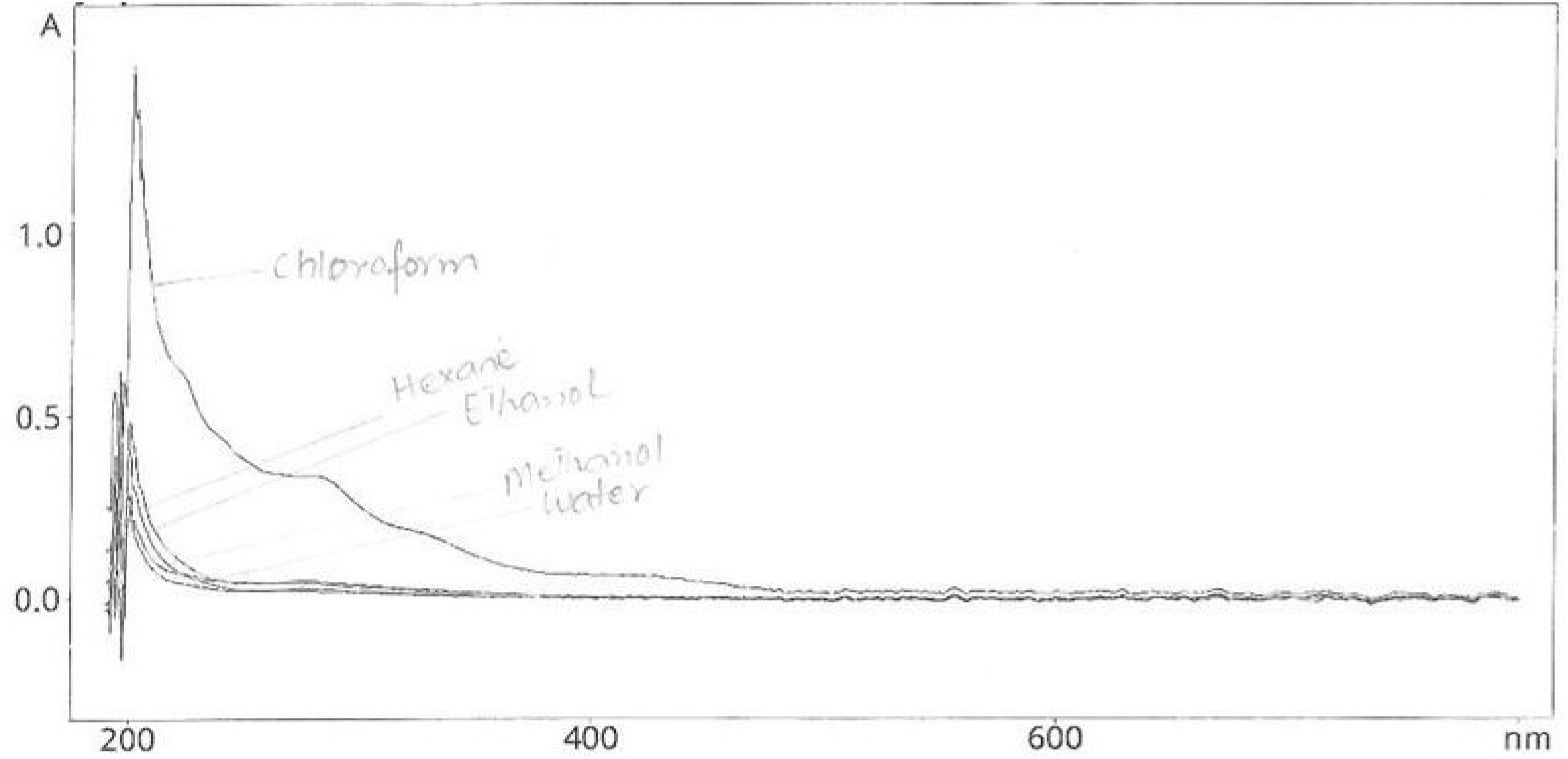
The results of extracts of *Beta vulagris* L. after UV-Vis analysis.

### 3.8 Analysis of hair tonic

Organoleptic examination of hair tonic showed that methanolic extract of 5% hair tonic solution had dark brown color while 10% had blackish brown color. The ethanolic extract of 5% tonic solution had light yellowish color while 10% had yellowish color. The concentration of tonic solution resulted in different colors because the more the solution was concentrated with extracts more deep color was produced. The odor of tonic was odorless. The pH of methanolic extract tonic solutions of 5% and 10% was 5.56 and 5.89 respectively and pH of ethanolic extract of tonic solutions of 5% and 10% was 6.01 and 6.24 respectively. The pH of the standard hair tonic that is minoxidil was 6.11.All of the hair tonics lie within the optimal pH range of skin which is 4.5-6.5 (Tranggono, 2007).

### 3.9 Activity of hair promotion

The application of different tonics on mice shown in figure 3, which were taken on day 7, day 14 and day 21. Totally there were five groups; group 1 have standard drug minoxidil as positive control, group 2 has no active pharmaceutical ingredient serve as negative group, group 3 was untreated group (no treatment was given), group 4 has methanolic extract hair tonic and group 5 has ethanolic extract hair tonic solution. In the given picture it can be clearly seen that both of the extracts had shown hair promoting activity.

**Figure 3.**
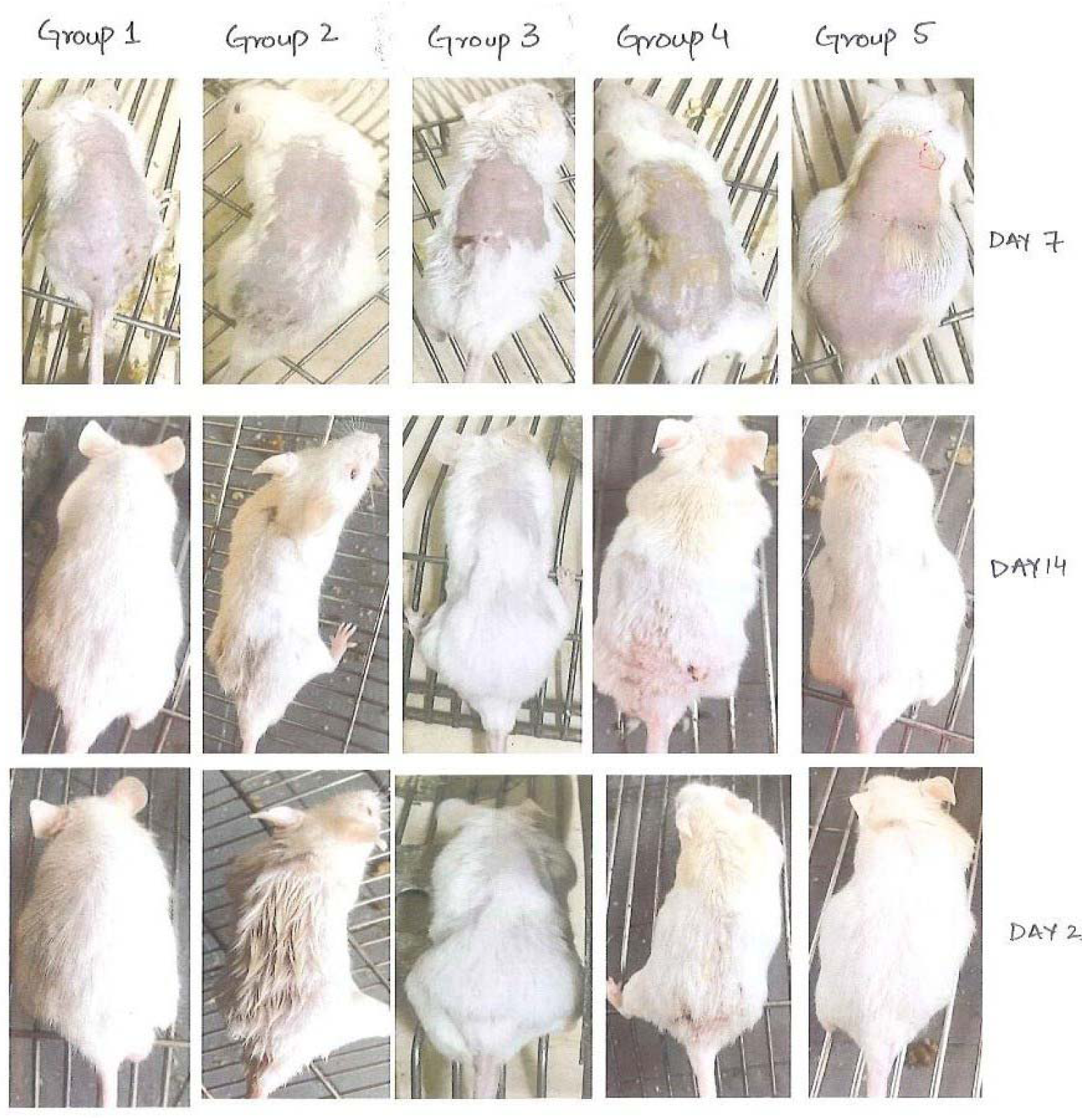
An application of different tonics on mice taken on day 7, day 14 and day 21 divided into five groups i.e. group 1 have standard drug minoxidil as positive control, group 2 has no active pharmaceutical ingredient serve as negative group, group 3 was untreated group, group 4 has methanolic extract hair tonic, and group 5 has ethanolic extract hair tonic solution.

### 3.10 Qualitative analysis of hair growth

Qualitative assessment of hair growth on the multiple groups has been shown in Table 7. The group one showed results on 15^th^ day, group two on 25^th^ day, group three on 30^th^ day, group four on 18^th^ day and group five on 17^th^ day.

**Table 7.** A qualitative assessment of hair growth.

| Groups | Qualitative evaluation (days) |
| --- | --- |
| Group 1 | 15 |
| Group 2 | 25 |
| Group 3 | 30 |
| Group 4 | 18 |
| Group 5 | 17 |

### 3.11 Length of the hair

The length of hair of mice was measured and given below in Table 8. In the first week of growth all the groups showed no activity, showing similar manners, while in the second week of growth, the positive group along with methanol, and ethanol group showed significant growth, but growth in ethanol and methanol group was less than positive group. The negative and untreated group showed growth but was not that significant as comparative to positive, methanol and ethanol. The third week, on day 21^st^ the positive group showed the maximum activity while methanol and ethanol showed the lesser activity. The methanol 5% and ethanol 5% showed activity which was slightly less than 10% solution. The 10 % methanol and ethanol tonic showed significant activity but was less as compared to standard group of minoxidil.

**Table 8.**
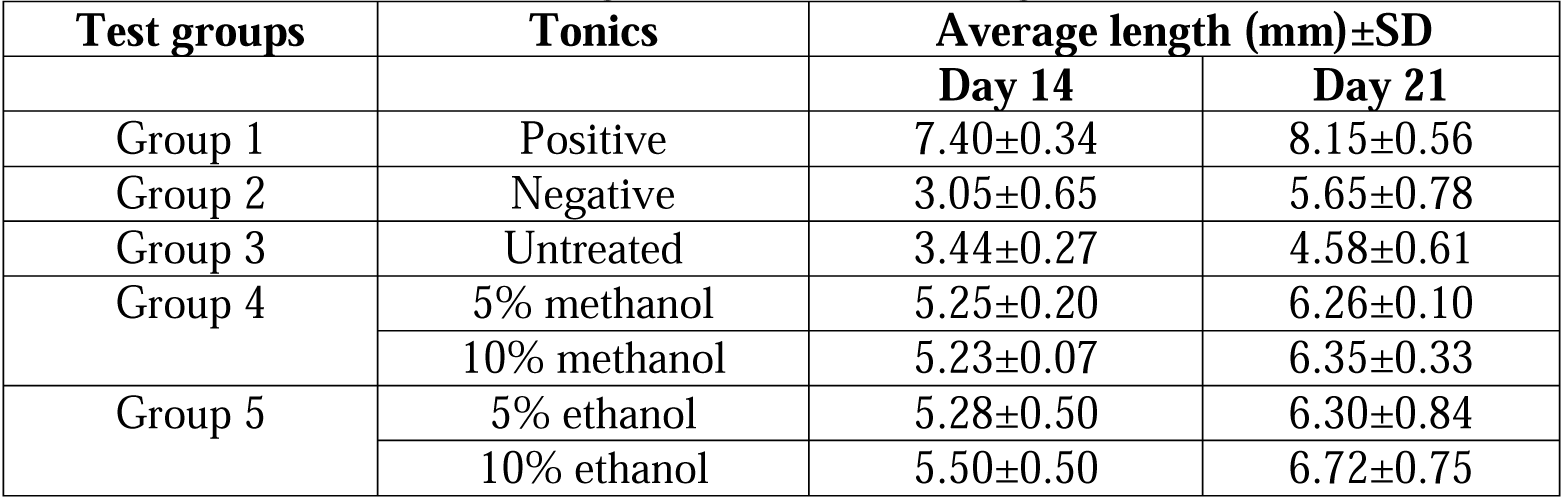
An estimation of length of hair of multiple groups.

| Test groups | Tonics | Average length (mm)±SD |  |
| --- | --- | --- | --- |
|  |  | Day 14 | Day 21 |
| Group 1 | Positive | 7.40±0.34 | 8.15±0.56 |
| Group 2 | Negative | 3.05±0.65 | 5.65±0.78 |
| Group 3 | Untreated | 3.44±0.27 | 4.58±0.61 |
| Group 4 | 5% methanol | 5.25±0.20 | 6.26±0.10 |
|  | 10% methanol | 5.23±0.07 | 6.35±0.33 |
| Group 5 | 5% ethanol | 5.28±0.50 | 6.30±0.84 |
|  | 10% ethanol | 5.50±0.50 | 6.72±0.75 |

The figure 4 given below showed the manner of activity in the different groups.

**Figure 4.**
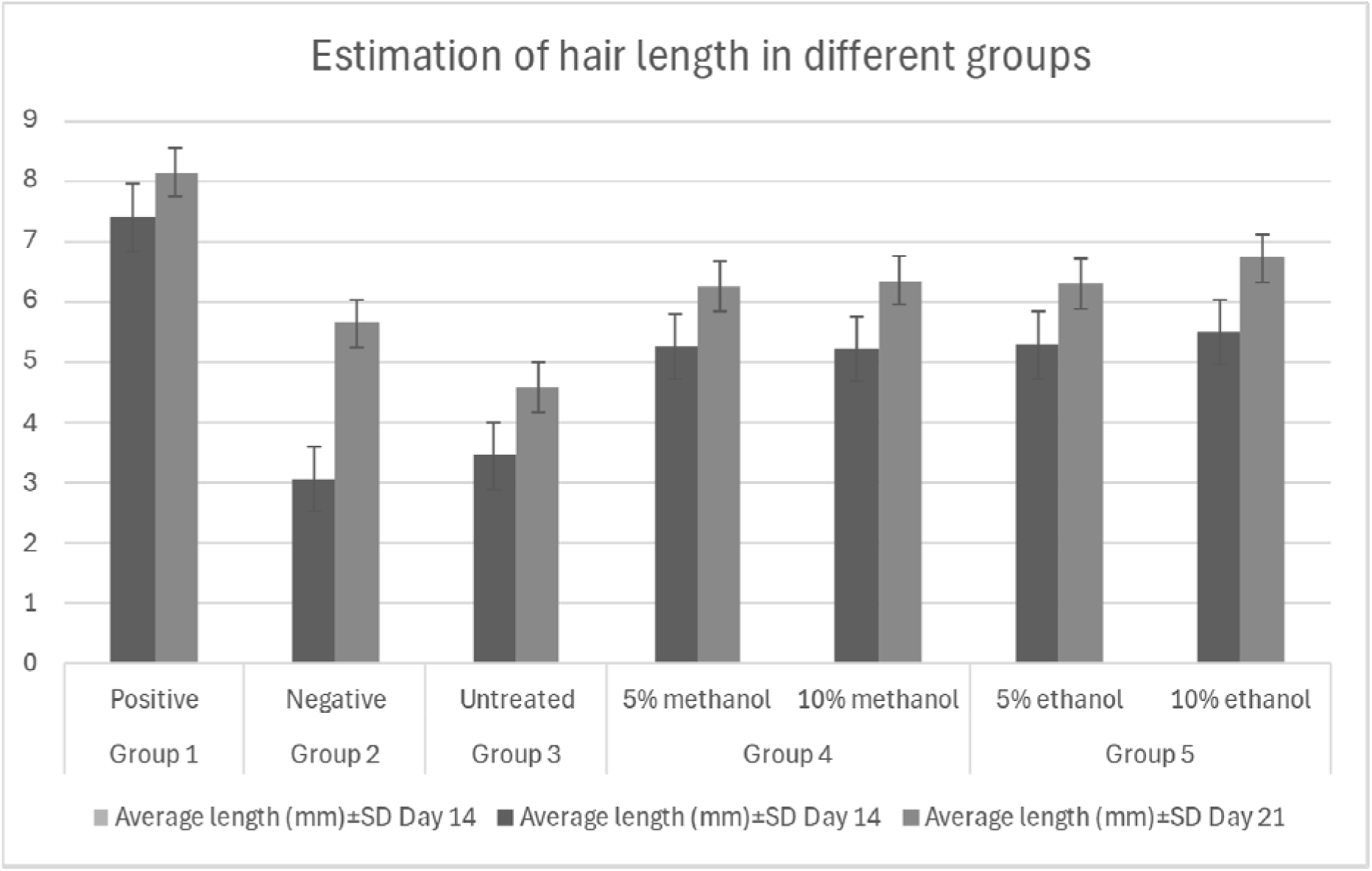
A comparison of all the groups’ length.

The statistical T-test was applied, and it showed that there was a significant difference p <0.05.

### 3.12 Measurement of hair diameter

The diameter of hair was measured, and the results are given in Table 9. The diameter of hair of different groups is shown in figure 5 below. When the group 4, methanol 5% and 10% wa compared with group 1(containing minoxidil), group 2 and group 3, it showed that positive group has greater activity in the second week and third week of growth while the methanol activity was less than positive group (minoxidil) but significant. Similarly, when group 5, ethanol 5% and 10% was compared with group 1, group 2 and group 3, it showed that all groups have the same activity except positive. But on the 21st day ethanol showed the significant increase in diameter but less than positive standard group containing minoxidil.

**Figure 5.**
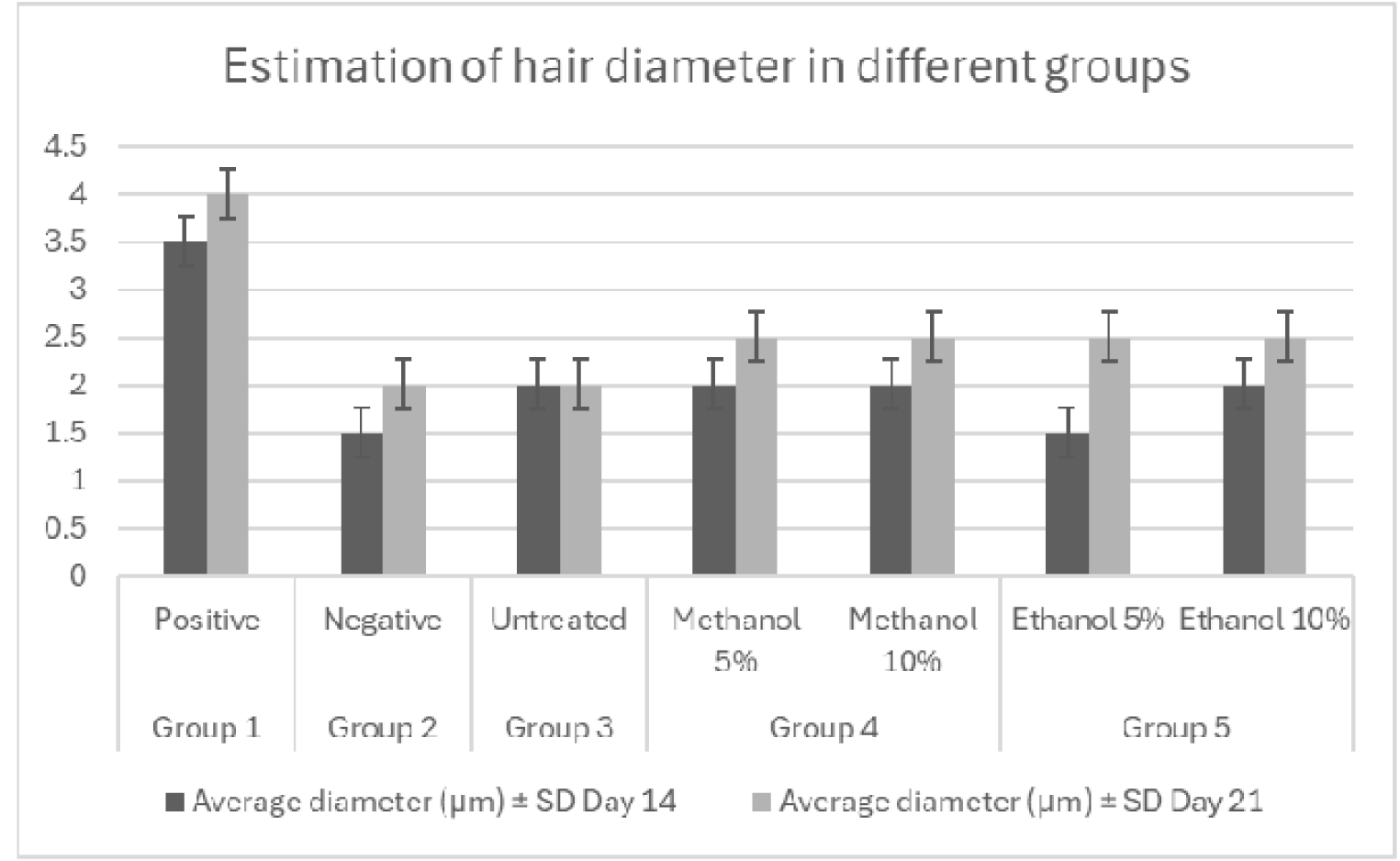
A comparison of all the groups’ diameter.

**Table 9.** The measurement of hair diameter.

| Test groups | Tonic | Average diameter ($\mu\text{m}$) ± SD | |
| --- | --- | --- | --- |
|  |  | Day 14 | Day 21 |
| Group 1 | Positive | 3.5±0.42 | 4±0.86 |
| Group 2 | Negative | 1.5±0.33 | 2±0.22 |
| Group 3 | Untreated | 2±0.34 | 2±0.24 |
| Group 4 | Methanol 5% | 2±0.41 | 2.5±0.65 |
| | Methanol 10% | 2±0.65 | 2.5±0.34 |
| Group 5 | Ethanol 5% | 1.5±0.67 | 2.5±0.42 |
| | Ethanol 10% | 2±0.88 | 2.5±0.54 |

The exact mechanism of increase in hair growth with the use of root of *Beta vulgaris* L. is not known. It might be due to the antioxidant property of the phenolic components, present in the root of *Beta vulgaris*, having strong reducing power, free scavenging radicals and hence increase cell proliferation (Ravichandran et al., 2012). The water-soluble nitrogenous components called betalains present in beetroot have powerful antioxidant and anti-inflammatory action, causing vasodilation to increase the blood flow to skin (Clifford et al., 2015; Lechner et al., 2010). Naturally many plants having anti-inflammatory action help in hair growth (Lourith and Kanlayavattanakul, 2013). Many nutrients, antioxidants and minerals cause an increase in hair growth due to multiple reasons (Rajendrasingh, 2018). Beet root has high potassium, which causes vasodilation hence to increase the blood flow and increases hair growth. Minoxidil is a vasodilator acting via ATP-potassium channels openers increasing potassium levels and promoting hair growth, but still exact mechanism is not known (Davies et al., 2005).It also possesses many other minerals but are not in significant quantities like sodium, calcium, zinc, iron but stills their presence is valuable. Minerals are helpful for the nourishment of healthy hair(Kaushik et al., 2011). The presence of saponins in the root of *Beta vulgaris* might be useful for hair growth. The exact phenomena are not known. But some of the herbs have shown that triterpenoids and saponins increase their hair growth (Chhikara et al., 2019; Lourith and Kanlayavattanakul, 2013; Matsuda et al., 2003).

## 4. Conclusion

All the values of physicochemical analysis were within range of given monograph. Total carbohydrates were highest than proteins and lipids in extracts of root of *Beta vulgaris* L. The different extracts were evaluated for secondary metabolites in phytochemical analysis. FTIR scan was performed and shows the presence of various functional groups present in extracts. By atomic absorption the mineral analysis was done, and it indicates high concentration of potassium and lower amount of sodium, calcium magnesium and phosphorous in the powder of *Beta vulgaris* L.UV-Vis analysis showed presence of conjugated systems and some polyphenolic compounds.

Hair promoting effect of root of *Beta vulgaris* was milder than the standard for both of the extracts but ethanol 5% and 10% showed more promoting result than methanol extracts. It can be used for hair growth purposes.

## Notes

### Competing Interest Statement

The authors declare that they have no known competing financial interests or personal relationships that could have appeared to influence the work reported in this paper

## References

Abubakar, M.A., Zulkifli, R.M., Hassan, W.N.A.W., Shariff, A.H.M., Malek, N.A.N.N., Zakaria, Z., Ahmad, F., 2015. Antibacterial properties of Persicaria minor (Huds.) ethanolic and aqueous-ethanolic leaf extracts. J. Appl. Pharm. Sci. 5, 050–056.

Adhirajan, N., Kumar, T.R., Shanmugasundaram, N., Babu, M., 2003. In vivo and in vitro evaluation of hair growth potential of Hibiscus rosa-sinensis Linn. J. Ethnopharmacol. 88, 235–239.

Aleixandre-Tudo, J.L., Nieuwoudt, H., Olivieri, A., Aleixandre, J.L., du Toit, W., 2018. Phenolic profiling of grapes, fermenting samples and wines using UV-Visible spectroscopy with chemometrics. Food Control 85, 11–22. 10.1016/j.foodcont.2017.09.014

Al-Hooti, S., Sidhu, J.S., Qabazard, H., 1998. Chemical composition of seeds of date fruit cultivars of United Arab Emirates.

Arawande, J.O., Akinnusotu, A., Alademeyin, J.O., 2018. Extractive Value and Phytochemical Screening of Ginger (zingiber officinale) and Turmeric (curcuma longa) Using Different Solvents.

Azwanida, N.N., 2015. A review on the extraction methods use in medicinal plants, principle, strength and limitation. Med Aromat Plants 4, 2167–0412.

Babarikins, D., Krūmiņa, G., Paegle, I., Amerika, D., Krūmiņa, Z., Vanags, D., Tihomirova, T., 2013. Allogeneic Bone Marrow Multipotent Mesenchymal Stromal Cells and Polytrauma Repair: The Role of Fractionated on the Basis of Molecular Mass Red Beetroot Juice in the Prevention of Transplanted Cells Side Effects in Rats. Proc. Latv. Acad. Sci. Sect. B Nat. Exact Appl. Sci. 67, 52–60. 10.2478/prolas-2013-0010

Babarykin, D., Smirnova, G., Pundinsh, I., Vasiljeva, S., Krumina, G., Agejchenko, V., 2019. Red Beet (Beta vulgaris) Impact on Human Health. J. Biosci. Med. 7, 61–79. 10.4236/jbm.2019.73007

Besbes, S., Blecker, C., Deroanne, C., Drira, N.-E., Attia, H., 2004. Date seeds: chemical composition and characteristic profiles of the lipid fraction. Food Chem. 84, 577–584.

Bezalwar Pratik, M., Gomashe Ashok, V., Gulhane Pranita, A., 2014. A quest of anti-acne potential of herbal medicines for extermination of MDR Staphylococcus aureus. Int. J. Pharm. Sci. Invent. 3, 12–17.

Caesar, L.K., Cech, N.B., 2019. Synergy and antagonism in natural product extracts: when 1 + 1 does not equal 2. Nat. Prod. Rep. 36, 869–888. 10.1039/C9NP00011A

Chaachouay, N., Zidane, L., 2024. Plant-Derived Natural Products: A Source for Drug Discovery and Development. Drugs Drug Candidates 3, 184–207. 10.3390/ddc3010011

Chanda, S., 2014. Importance of pharmacognostic study of medicinal plants: An overview. J. Pharmacogn. Phytochem. 2.

Che, C.-T., George, V., Ijinu, T.P., Pushpangadan, P., Andrae-Marobela, K., 2024. Chapter 2 - Traditional medicine, in: McCreath, S.B., Clement, Y.N. (Eds.), Pharmacognosy (Second Edition). Academic Press, pp. 11–28. 10.1016/B978-0-443-18657-8.00037-2

Chen, G., Liu, S., Chen, S., Qi, Z., 2001. FTIR Spectra, Thermal Properties, and Dispersibility of a Polystyrene/Montmorillonite Nanocomposite. Macromol. Chem. Phys. 202, 1189–1193. 10.1002/1521-3935(20010401)202:7<1189::AID-MACP1189>3.0.CO;2-M

Chhikara, N., Kushwaha, K., Sharma, P., Gat, Y., Panghal, A., 2019. Bioactive compounds of beetroot and utilization in food processing industry: A critical review. Food Chem. 272, 192–200.

Claro, A.E., Palanza, C., Mazza, M., Schuenemann, G.E.U.M., Rigoni, M., Pontecorvi, A., Janiri, L., Pitocco, D., Muti, P., 2024. Historical use of medicinal plants and future potential from phytotherapy to phitochemicals. Ann. Bot. 14. 10.13133/2239-3129/18564

Clifford, T., Howatson, G., West, D.J., Stevenson, E.J., 2015. The potential benefits of red beetroot supplementation in health and disease. Nutrients 7, 2801–2822.

Davies, G.C., Thornton, M.J., Jenner, T.J., Chen, Y.-J., Hansen, J.B., Carr, R.D., Randall, V.A., 2005. Novel and established potassium channel openers stimulate hair growth in vitro: implications for their modes of action in hair follicles. J. Invest. Dermatol. 124, 686–694.

Dhami, L., 2021. Psychology of Hair Loss Patients and Importance of Counseling. Indian J. Plast. Surg. 54, 411–415. 10.1055/s-0041-1741037

Dhawan, D., Gupta, J., 2017. Research article comparison of different solvents for phytochemical extraction potential from datura metel plant leaves. Int J Biol Chem 11, 17–22.

El Gamal, A.A., AlSaid, M.S., Raish, M., Al-Sohaibani, M., Al-Massarani, S.M., Ahmad, A., Hefnawy, M., Al-Yahya, M., Basoudan, O.A., Rafatullah, S., 2014. Beetroot ( *Beta vulgaris* L.) Extract Ameliorates Gentamicin-Induced Nephrotoxicity Associated Oxidative Stress, Inflammation, and Apoptosis in Rodent Model. Mediators Inflamm. 2014, 1–12. 10.1155/2014/983952

El-Hendawy, A.-N.A., 2006. Variation in the FTIR spectra of a biomass under impregnation, carbonization and oxidation conditions. J. Anal. Appl. Pyrolysis 75, 159–166.

Essiett, U.A., Bala, D., 2011. Phytochemical and physicochemical analysis of the leaves of Laportea aestuans (Linn.) Chew and Laportea ovalifolia (Schumach.) Chew (male and female). Asian J. Plant Sci. Res.

Guiné, R.P.F., A. M. Castro, J.A., 2003. Analysis of Moisture Content and Density of Pears During Drying. Dry. Technol. 21, 581–591. 10.1081/DRT-120018464

Imtiaz, F., Islam, M., Saeed, H., Saleem, B., Asghar, M., Saleem, Z., 2017. Impact of Trigonella foenum-graecum leaves extract on mice hair growth. Pak. J. Zool. 49.

Ji, S., Zhu, Z., Sun, X., Fu, X., 2021. Functional hair follicle regeneration: an updated review. Signal Transduct. Target. Ther. 6, 66.

Julkunen-Tiitto, R., Nenadis, N., Neugart, S., Robson, M., Agati, G., Vepsäläinen, J., Zipoli, G., Nybakken, L., Winkler, B., Jansen, M.A.K., 2015. Assessing the response of plant flavonoids to UV radiation: an overview of appropriate techniques. Phytochem. Rev. 14, 273–297. 10.1007/s11101-014-9362-4

Kapadia, G.J., Rao, G.S., 2013. Anticancer Effects of Red Beet Pigments, in: Neelwarne, B. (Ed.), Red Beet Biotechnology. Springer US, Boston, MA, pp. 125–154. 10.1007/978-1-4614-3458-0_7

Kaushik, R., Gupta, D., Yadav, R., 2011. Alopecia: herbal remedies. Int. J. Pharm. Sci. Res. 2, 1631.

Klevay, L.M., Combs Jr, G.F., 2005. Mineral elements related to cardiovascular health. Nutr. Drink. Water 92.

Kumar, S., Brooks, M.S.-L., 2018. Use of Red Beet (Beta vulgaris L.) for Antimicrobial Applications—a Critical Review. Food Bioprocess Technol. 11, 17–42. 10.1007/s11947-017-1942-z

Lechner, J.F., Wang, L.-S., Rocha, C.M., Larue, B., Henry, C., McIntyre, C.M., Riedl, K.M., Schwartz, S.J., Stoner, G.D., 2010. Drinking Water with Red Beetroot Food Color Antagonizes Esophageal Carcinogenesis in *N*-Nitrosomethylbenzylamine-Treated Rats. J. Med. Food 13, 733–739. 10.1089/jmf.2008.0280

Lemarchand, C., Gref, R., Couvreur, P., 2004. Polysaccharide-decorated nanoparticles. Eur. J. Pharm. Biopharm. 58, 327–341.

Li, X., Wang, X., Wang, C., Zhang, J., Zhou, C., 2022. Hair Shedding Evaluation for Alopecia: A Refined Wash Test. Clin. Cosmet. Investig. Dermatol. 15, 117–126. 10.2147/CCID.S347898

Liu, C., 2021. Overview on development of ASEAN traditional and herbal medicines. Chin. Herb. Med., RCEP Traditional Medicine Research 13, 441–450. 10.1016/j.chmed.2021.09.002

Lourith, N., Kanlayavattanakul, M., 2013. Hair loss and herbs for treatment. J. Cosmet. Dermatol. 12, 210–222. 10.1111/jocd.12051

Lowry, O.H., Rosebrough, N.J., Farr, A.L., Randall, R.J., 1951. Protein measurement with the Folin phenol reagent.

Maheshwari, R.K., Parmar, V., Joseph, L., 2013. Latent therapeutic gains of beetroot juice. World J. Pharm. Res. 2, 804–820.

Manohar, C.M., Kundgar, S.D., Doble, M., 2017. Betanin immobilized LDPE as antimicrobial food wrapper. LWT 80, 131–135. 10.1016/j.lwt.2016.07.020

Martinez, R.M., Longhi-Balbinot, D.T., Zarpelon, A.C., Staurengo-Ferrari, L., Baracat, M.M., Georgetti, S.R., Sassonia, R.C., Verri, W.A., Casagrande, R., 2015. Anti-inflammatory activity of betalain-rich dye of Beta vulgaris: effect on edema, leukocyte recruitment, superoxide anion and cytokine production. Arch. Pharm. Res. 38, 494–504. 10.1007/s12272-014-0473-7

Matsuda, H., Yamazaki, M., Asanuma, Y., Kubo, M., 2003. Promotion of hair growth by ginseng radix on cultured mouse vibrissal hair follicles. Phytother. Res. 17, 797–800. 10.1002/ptr.1241

Meda, A., Lamien, C.E., Romito, M., Millogo, J., Nacoulma, O.G., 2005. Determination of the total phenolic, flavonoid and proline contents in Burkina Fasan honey, as well as their radical scavenging activity. Food Chem. 91, 571–577.

Miraj, S., 2016. Chemistry and pharmacological effect of beta vulgaris: A systematic review. Pharm. Lett. 8, 404–409.

Mir-Marques, A., Cervera, M.L., de la Guardia, M., 2016. Mineral analysis of human diets by spectrometry methods. TrAC Trends Anal. Chem. 82, 457–467.

Mirmiran, P., Houshialsadat, Z., Gaeini, Z., Bahadoran, Z., Azizi, F., 2020. Functional properties of beetroot (Beta vulgaris) in management of cardio-metabolic diseases. Nutr. Metab. 17, 3. 10.1186/s12986-019-0421-0

Mohammed, A.M., 2018. UV-Visible Spectrophotometric Method and Validation of Organic Compounds. Eur. J. Eng. Technol. Res. 3, 8–11. 10.24018/ejeng.2018.3.3.622

Nandaniya, H., Khanpara, P., Faldu, D.S., 2023. Focus on herbal home remedies for hair regrowth and loss. J. Pharmacogn. Phytochem. 12, 327–340. 10.22271/phyto.2023.v12.i5d.14744

Natarelli, N., Gahoonia, N., Sivamani, R.K., 2023. Integrative and Mechanistic Approach to the Hair Growth Cycle and Hair Loss. J. Clin. Med. 12, 893. 10.3390/jcm12030893

Object, object, n.d. Phytochemical, Proximate and Mineral Analyses of the Leaves of Gossypium hirsutum L. and Momordica charantia L.

Ovchinnikov, O.V., Evtukhova, A.V., Kondratenko, T.S., Smirnov, M.S., Khokhlov, V.Y., Erina, O.V., 2016. Manifestation of intermolecular interactions in FTIR spectra of methylene blue molecules. Vib. Spectrosc. 86, 181–189.

Pallab, K., Tapan, B., Tapas, P., Ramen, K., 2013. Estimation of total flavonoids content (TPC) and antioxidant activities of methanolic whole plant extract of Biophytum sensitivum Linn. J. Drug Deliv. Ther. 3, 33–37.

Pan, S.-Y., Litscher, G., Gao, S.-H., Zhou, S.-F., Yu, Z.-L., Chen, H.-Q., Zhang, S.-F., Tang, M.-K., Sun, J.-N., Ko, K.-M., 2014. Historical Perspective of Traditional Indigenous Medical Practices: The Current Renaissance and Conservation of Herbal Resources. Evid. Based Complement. Alternat. Med. 2014, 525340. 10.1155/2014/525340

Petit, T., Puskar, L., 2018. FTIR spectroscopy of nanodiamonds: Methods and interpretation. Diam. Relat. Mater. 89, 52–66.

Petrovska, B.B., 2012. Historical review of medicinal plants’ usage. Pharmacogn. Rev. 6, 1–5. 10.4103/0973-7847.95849

Rajendrasingh, R.R., 2018. Nutritional Correction for Hair Loss, Thinning of Hair, and Achieving New Hair Regrowth, in: Pathomvanich, D., Imagawa, K. (Eds.), Practical Aspects of Hair Transplantation in Asians. Springer Japan, Tokyo, pp. 667–685. 10.1007/978-4-431-56547-5_71

Ravichandran, K., Ahmed, A.R., Knorr, D., Smetanska, I., 2012. The effect of different processing methods on phenolic acid content and antioxidant activity of red beet. Food Res. Int. 48, 16–20.

Rop, O., Mlcek, J., Jurikova, T., Neugebauerova, J., Vabkova, J., 2012. Edible flowers—a new promising source of mineral elements in human nutrition. Molecules 17, 6672–6683.

Rose, M.H., Sudha, P., Sudhakar, K., 2014. Effect of antioxidants and hepatoprotective activities of methanol extract of beet root (Beta vulgaris L.) against carbon tetrachloride induced hepatotoxicity in rat models. Int J Pharm Sci Res 5, 2546.

Sabarwal, N., Varghese, D., Barik, R., Khandelwal, A., Jain, A.J.S., Jain, S., 2009. Development and evaluation of polyherbal formulations for hair growth activity. PharmacogNet 1, 165–70.

Saleem, U., Hussain, K., Ahmad, M., Irfan Bukhari, N., Malik, A., Ahmad, B., 2014. Physicochemical and phytochemical analysis of Euphorbia helioscopia (L.). Pak. J. Pharm. Sci. 27.

Scalbert, A., Johnson, I.T., Saltmarsh, M., 2005. Polyphenols: antioxidants and beyond. Am. J. Clin. Nutr. 81, 215S–217S.

Shi, J., Arunasalam, K., Yeung, D., Kakuda, Y., Mittal, G., Jiang, Y., 2004. Saponins from Edible Legumes: Chemistry, Processing, and Health Benefits. J. Med. Food 7, 67–78. 10.1089/109662004322984734

Sim, C.O., Hamdan, M.R., Ismail, Z., Ahmad, M.N., 2004. Assessment of herbal medicines by chemometrics–assisted interpretation of FTIR spectra. J Anal. Chim. Acta 1, 14.

Singh, B., Hathan, B.S., 2014. Composição química, propriedades funcionais e processamento de beterraba-uma revisão. Int J Sci Eng Res 5, 679–684.

Sisa, M., Bonnet, S.L., Ferreira, D., Van der Westhuizen, J.H., 2010. Photochemistry of flavonoids. Molecules 15, 5196–5245.

Soetan, K.O., Olaiya, C.O., Oyewole, O.E., 2010. The importance of mineral elements for humans, domestic animals and plants: A review. Afr. J. Food Sci. 4, 200–222.

Sparg, S., Light, M.E., Van Staden, J., 2004. Biological activities and distribution of plant saponins. J. Ethnopharmacol. 94, 219–243.

Tranggono, R.I., 2007. BP: Ilmu Pengetahuan Kosmetik. Gramedia Pustaka Utama.

Willcox, M.L., Bodeker, G., 2004. Traditional herbal medicines for malaria. Bmj 329, 1156–1159.

Yadav, R.N.S., Agarwala, M., 2011. Phytochemical analysis of some medicinal plants. J. Phytol. 3.

Zagatto, E., Worsfold, P.J., 2017. Spectrophotometry: Overview. Reference Module in Chemistry, Molecular Sciences and Chemical Engineering.

Zamberlin, Š., Antunac, N., Havranek, J., Samaržija, D., 2012. Mineral elements in milk and dairy products. Mljekarstvo Časopis Za Unaprjedjenje Proizv. Prerade Mlijeka 62, 111–125.

